# The structure of a highly conserved picocyanobacterial protein reveals a Tudor domain with an RNA binding function

**DOI:** 10.1101/542068

**Authors:** Katherine M Bauer, Rose Dicovitsky, Maria Pellegrini, Olga Zhaxybayeva, Michael J Ragusa

## Abstract

Cyanobacteria of the *Prochlorococcus* and marine *Synechococcus* genera are the most abundant photosynthetic microbes in the ocean. Intriguingly, the genomes of these bacteria are very divergent even within each genus, both in gene content and at the amino acid level of the encoded proteins. One striking exception to this is a 62 amino acid protein, termed *<u>P</u>rochlorococcus*/*<u>S</u>ynechococcus* <u>H</u>yper <u>C</u>onserved <u>P</u>rotein (PSHCP). PSHCP is not only found in all sequenced *Prochlorococcus* and marine *Synechococcus* genomes but it is also nearly 100% identical in its amino acid sequence across all sampled genomes. Such universal distribution and sequence conservation suggests an essential cellular role of the encoded protein in these bacteria. However, the function of PSHCP is unknown. We used Nuclear Magnetic Resonance (NMR) spectroscopy to determine its structure. We found that 53 of the 62 amino acids in PSHCP form a Tudor domain, while the remainder of the protein is disordered. NMR titration experiments revealed that PSHCP has only a weak affinity for DNA, but an 18.5-fold higher affinity for tRNA, hinting at an involvement of PSHCP in translation. Isothermal titration calorimetry experiments further revealed that PSHCP also binds single-stranded, double-stranded and hairpin RNAs. These results provide the first insight into the structure and function of PSHCP, demonstrating that PSHCP is an RNA binding protein that can recognize a broad array of RNA molecules.

---

With the mean annual average of 3.6 × 10^27^ cells, cyanobacteria from genera *Prochlorococcus* (1) and *Synechococcus* (2) numerically dominate microbial communities of the global ocean (3). Because of such abundance, these photosynthesizing bacteria are extremely important players in the global carbon cycle (3). The small genomes and limited gene content of individual *Prochlorococcus* and *Synechococcus* cells (1.6-2.9 Mb, encoding ~1700-3100 genes; (4)) are counterbalanced by a diverse collective gene pool, which in the case of *Prochlorococcus* is estimated to contain 80,000 genes (5,6). The genomes of *Prochlorococcus* and *Synechococcus* are also surprisingly divergent: in pairwise comparisons, the genome-wide average amino acid identity (AAI; (7)) of the encoded proteins is often below 60% even within each genus (8). Maintenance of such remarkable sequence and gene content divergence remains poorly understood (6). However, one protein-coding gene is a curious exception, with its product showing almost 100% amino acid sequence conservation across all currently available *Prochlorococcus* and marine *Synechococcus* genomes ((9,10) and Supplementary Datasets 1 and 2). This gene encodes a 62 amino acid protein of unknown function, dubbed PSHCP for <u>*P*</u>*rochlorococcus*/<u>*S*</u>*ynechococcus* <u>H</u>yper <u>C</u>onserved <u>P</u>rotein, (9,10). Although at the time of its discovery the gene encoding PSHCP was limited to the cyanobacterial clade consisting of *Prochlorococcus* and marine *Synechococcus*, sequence similarity searches now identify the PSHCP gene in genomes of freshwater *Synechococcus* (11); several *Cyanobium* spp. from both freshwater and marine environments; in cyanobacterial sponge symbiont “Candidatus *Synechococcus spongarium*” (12); in three species of *Paulinella*, which is a photosynthetic protist with a chromatophore thought to be derived from marine *Synechococcus* spp. (13); and in metagenomically-assembled cyanobacteria from globally distributed marine, brackish and freshwater environments. Within cyanobacteria, all characterized PSHCP-containing organisms form a clade (14) within the order *Synechococcales* (15). Hence, the PSHCP protein remains extremely conserved in a specific subgroup of cyanobacteria (Supplementary Datasets 1 and 2) but is undetectable outside of this subgroup.

The remarkable conservation of the PSHCP protein and retention of this gene even within extremely reduced chromatophore genomes (13) suggest that this protein may encode an important housekeeping function. In three examined strains of *Prochlorococcus* and marine *Synechococcus,* the gene is expressed, and its protein product is abundant (10). A large proportion of positively charged amino acids (16%) (9), a predicted isoelectric point of 11.3 (9), and the protein’s association with the ribosomal protein L2 in pull-down assays (10) prompted the hypotheses of PSHCP involvement in binding of either DNA or RNA, and its possible association with a ribosome. Yet, the amino acid sequence of PSHCP shows no significant sequence similarity to any protein domain with a known function.

To gain more insight into the possible function of PSHCP, we determined the structure of the PSHCP protein using Nuclear Magnetic Resonance (NMR) spectroscopy and investigated the protein’s interactions with nucleic acids *in vitro*.

## RESULTS

### The structure of PSHCP

We began by screening different constructs of PSHCP for their behavior for structure determination. An initial round of construct screening revealed that residues 57-62 could be removed without altering the overall stability of the protein. This is in good agreement with disorder prediction using the IUPred server (16), which suggests that residues 57-62 are disordered. PSHCP_1-56_ could be easily concentrated above 1 mM making it ideal for structure determination. Secondary structure probabilities as calculated using TALOS-N (17) from ^1^H,^13^C,^15^N chemical shifts, show that the first three residues of PSHCP_1-56_ are disordered and that the remainder of the protein contains five β-strands (Fig. S1A). To further investigate the dynamics of PSHCP_1-56_ we performed a {^1^H}-^15^N heteronuclear Nuclear Overhauser Effect (NOE) experiment. In the {^1^H}-^15^N heteronuclear NOE experiment a peak intensity ratio closer to one indicates little motion of the N-H bond on the picosecond to nanosecond timescale while a smaller peak intensity ratio corresponds to more motion on this time scale and thus residues which are disordered. The {^1^H}-^15^N heteronuclear NOE data recorded on PSHCP_1-56_ is in good agreement with the secondary structure probabilities confirming that the structured core of the protein corresponds to residues 4-56 (Fig. S1B).

For the structure calculation of PSHCP_1-56_, 1195 NOE derived distance constraints and 44 dihedral angle constraints were used (Table 1). Excluding the first three amino acids, this resulted in 22.1 distance constraints per amino acid. The 20 lowest energy structures align with an average pairwise backbone RMSD of 0.72 ± 0.15 Å for residues 4-29 and 35-56 (Fig. 1A). Residues 30-34 form a flexible loop between β-strands two and three, and thus were excluded from the RMSD calculation. The overall fold of PSHCP contains five β-strands, which form a single antiparallel β-sheet with an overall barrel-like architecture (Fig. 1B and 1C).

**Table 1.**
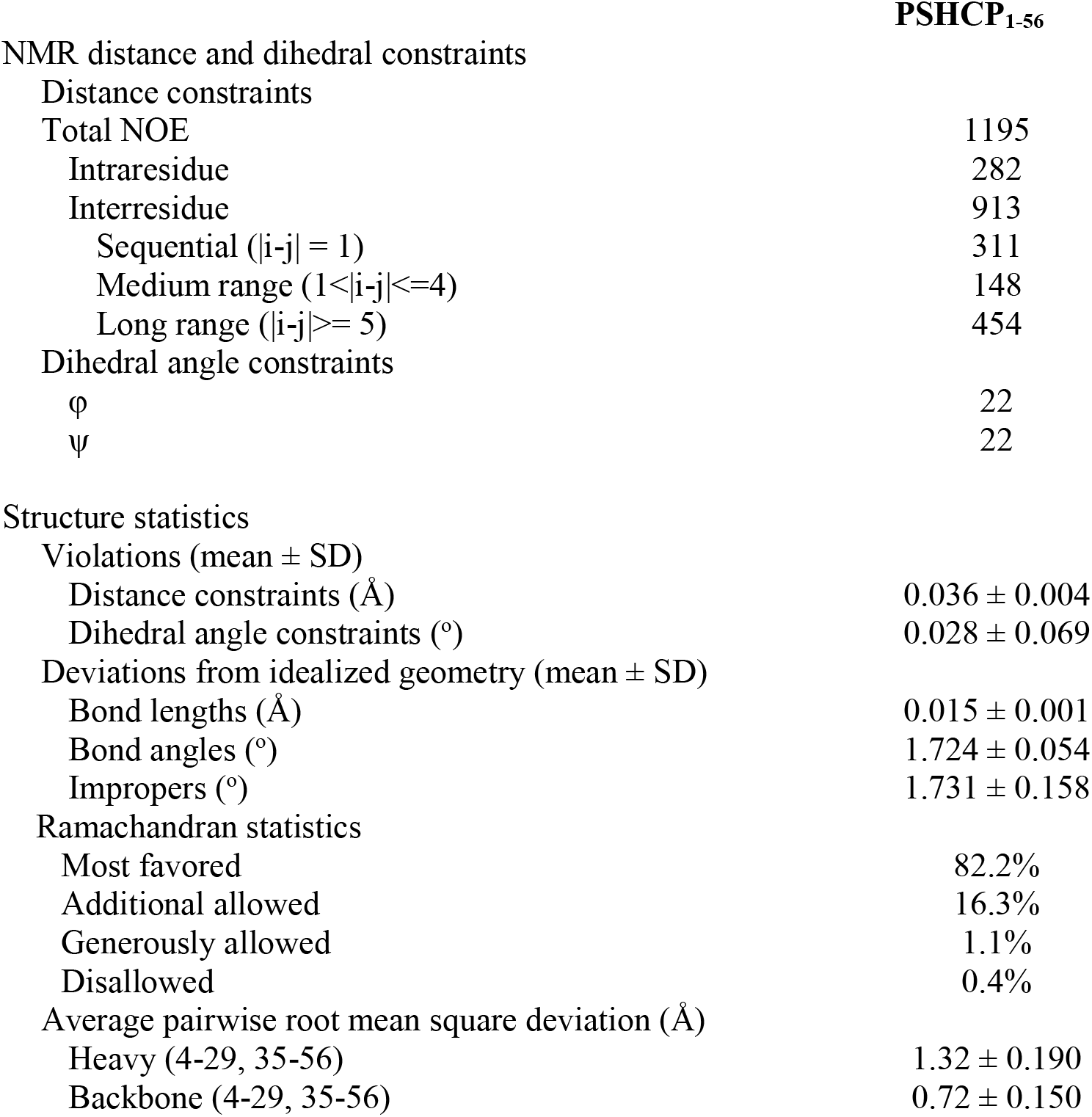
Structural statistics of PSHCP_1-56_

**Figure 1.**
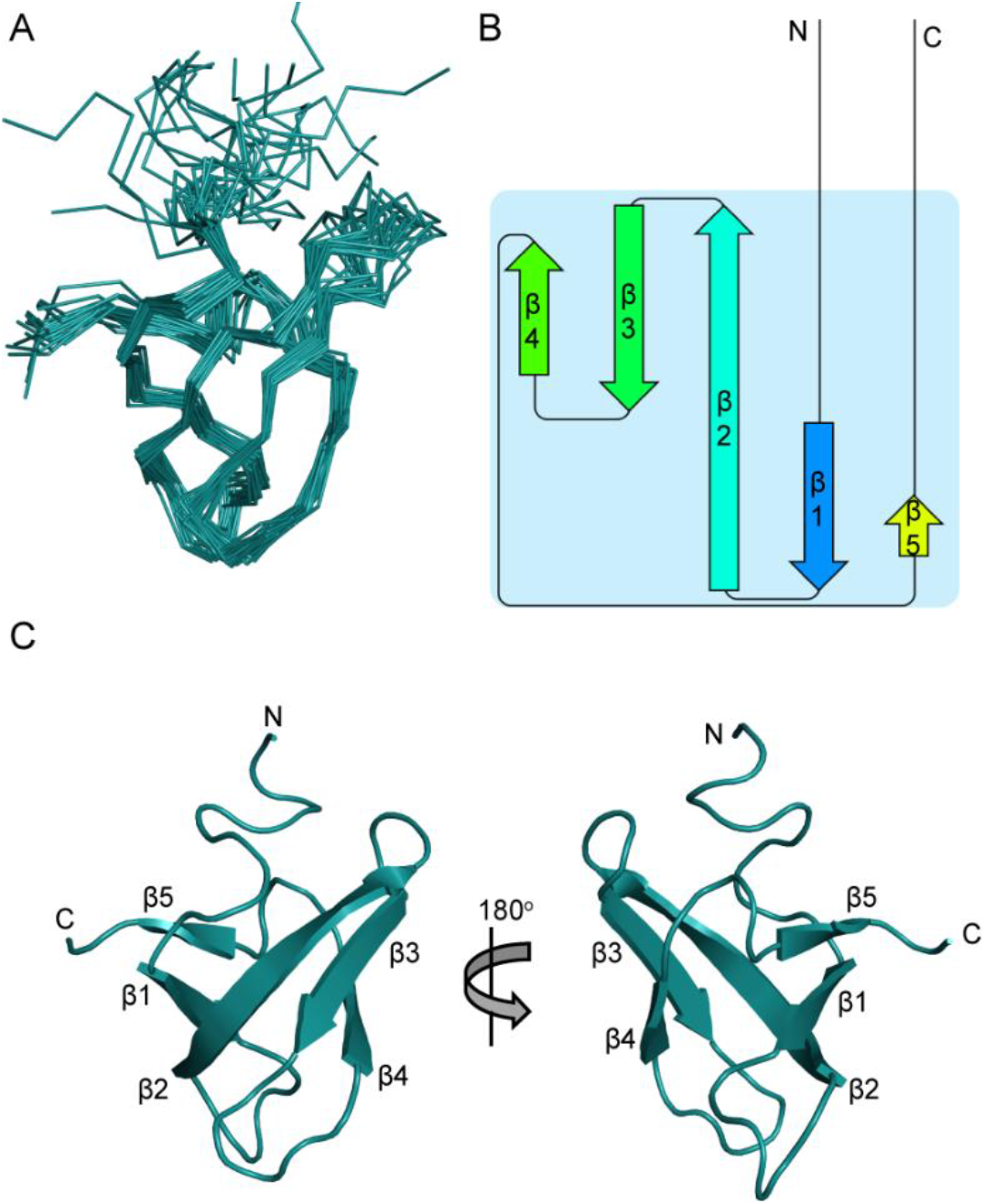
The structure of PSHCP. A) Ribbon diagram of PSHCP_1-56_ showing the overlay of the 20 lowest energy structures. B) Topology diagram of PSHCP generated using the pro-origami webserver (48). C) Cartoon representation of PSHCP shown in two different orientations. Each β-strand in the structure is labeled and numbered. The N and C termini are also labeled.

To determine if PSHCP is part of a conserved domain family, we used the Dali server (18) to search the Protein Data Bank (PDB) for proteins with structural similarity to PSHCP. PSHCP aligns well to a number of Tudor domains including the Tudor domain from SMN (Fig. S2) (19). The highest structural similarity is to the Tudor domain from the human TDRD3 protein: the 53 amino acids of PSHCP (residues 4-56) align to it with a backbone RMSD of 1.5 Å (Fig. 2A-C). Tudor domains are small, approximately 60 amino acid long, domains containing four or five β-strands in a barrel-like fold. Most known Tudor domains are found in eukaryotes (98.97% of the InterPro records for Tudor domain), where they are often present in 2-3 adjacent copies within large, multidomain proteins (20). In contrast, full length PSHCP consists entirely of a single Tudor domain (residues 4-56).

**Figure 2.**
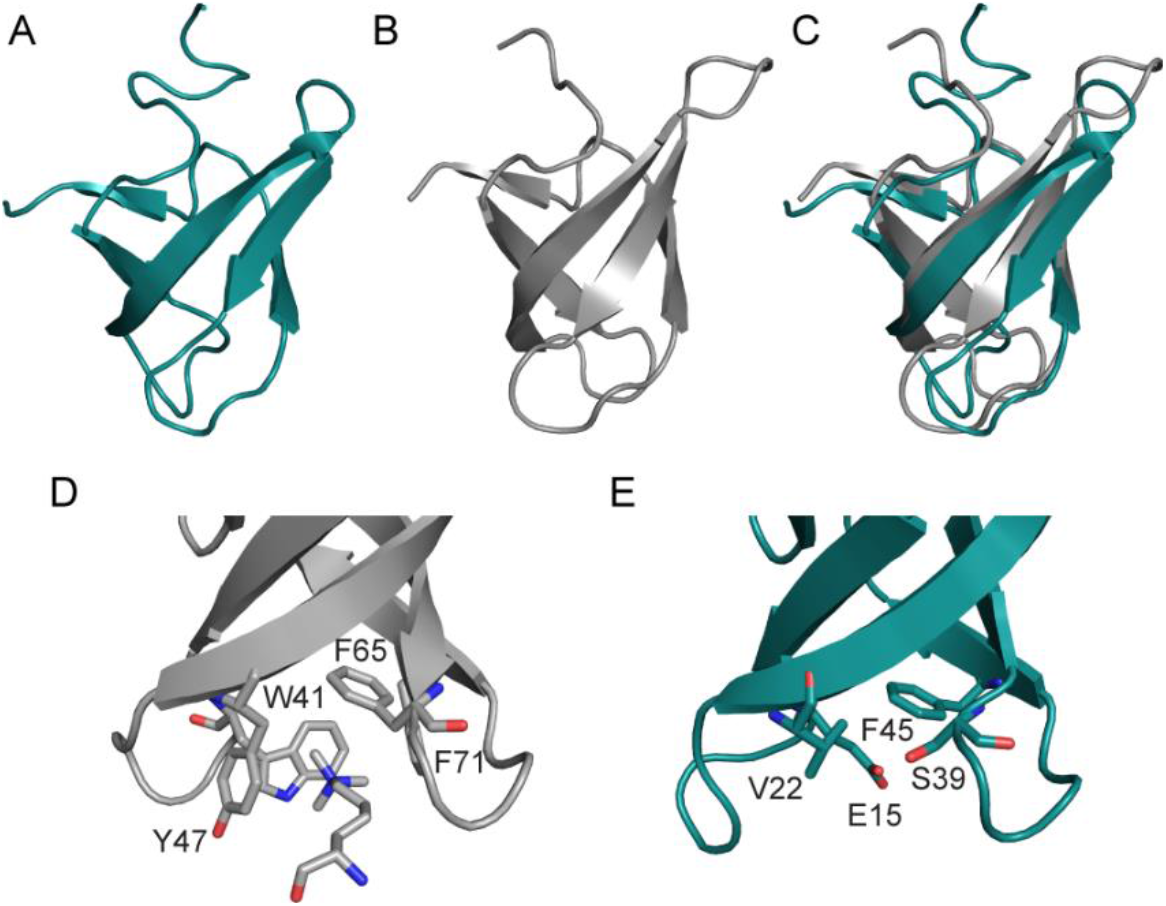
PSHCP contains a Tudor domain. A) Cartoon representation of PSHCP. B) Cartoon representation of the Tudor domain from human Tudor domain-containing protein 3 (PDBID: 3PMT). C) Overlay of the structures from panels A and B. D) A close-up view of the aromatic cage from the Tudor domain of PHF1 (PDBID: 4HCZ) bound to trimethylated lysine. E) A close-up view showing the residues in PSHCP that are located in the position of the aromatic cage residues in typical Tudor domains. In panels D and E all residues that make up the aromatic cage are shown as stick representations and labeled.

### PSHCP is unlikely to be involved in recognition of methylated proteins

The best-characterized functions of Tudor domains are to bind methylated lysine and arginine residues (20,21). This frequently results in the targeting of Tudor domain containing proteins to histones to regulate DNA damage responses and chromatin remodeling. The interaction of Tudor domains with methylated residues occurs through a conserved aromatic cage (Fig. 2D) (20,22). This aromatic cage is composed of four aromatic residues, which are individually located in β-strands one through four of the Tudor domain. However, PSHCP contains only one aromatic residue in any of the positions corresponding to the aromatic cage residues and is therefore unlikely to bind methylated amino acids (Fig. 2E).

### PSHCP has a weak affinity for DNA

Tudor domains have also been shown to regulate DNA function by directly binding DNA (23,24). In PSHCP, ten of 62 amino acids are positively charged (Fig. 3A). The majority of these residues cluster in one of two locations on the surface of PSHCP, generating two distinct positively charged patches that may support DNA binding (Fig. 3B). To determine if PSHCP is able to bind to DNA, we incubated ^15^N-labeled PSHCP_1-56_ with different concentrations of a double-stranded GC-rich DNA sequence (dsGCDNA) that has been previously shown to bind the double interdigitated Tudor domain of retinoblastoma-binding protein 1 (RBBP1) (23). When titrating dsGCDNA into PSHCP seven peaks shifted in the PSHCP_1-56_ ^1^H-^15^N HSQC spectrum, demonstrating that PSHCP_1-56_ can bind to DNA (Fig. 4A). As only seven peaks shifted upon DNA binding, this suggests that PSHCP_1-56_ does not undergo large structural changes upon DNA binding. The peaks for L14, E15, S16, G20 and V22 all shifted upon DNA binding. These five residues all sit in the loop between β-strand 1 and 2 suggesting that this region is important for DNA binding.

**Figure 3.**
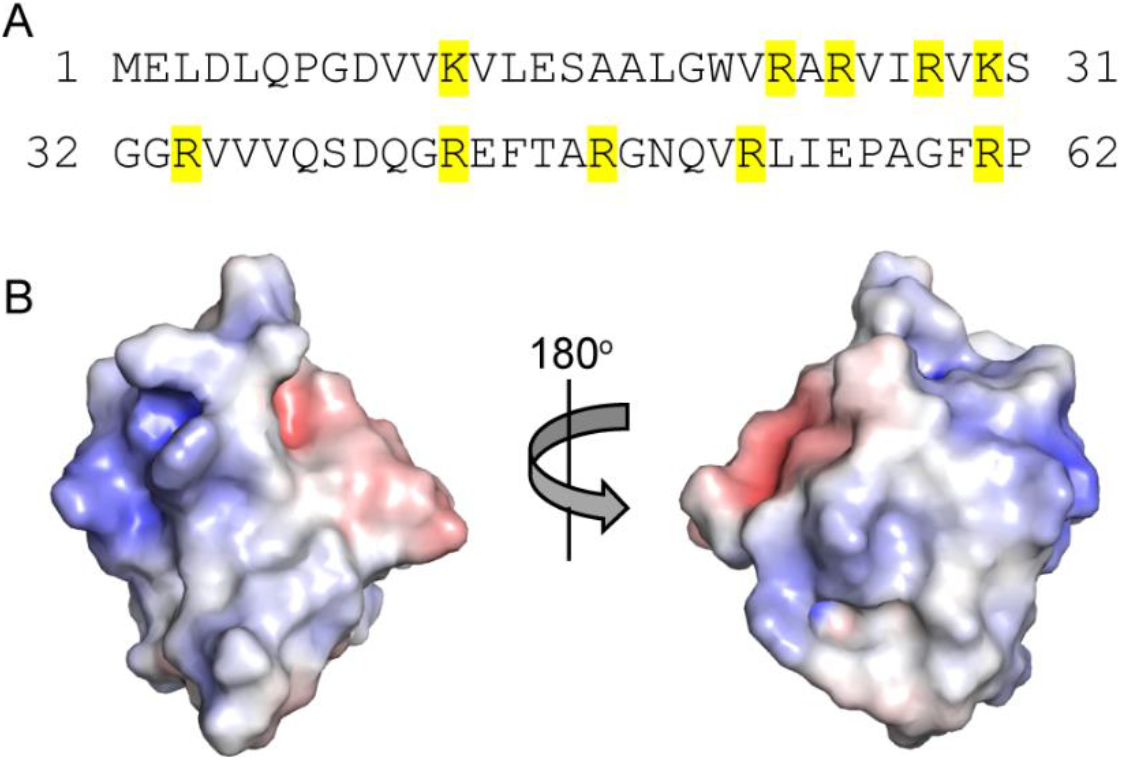
Electrostatics of PSHCP. A) The amino acid sequence of the PSHCP from *P. marinus* strain CCMP1375. All positively charge residues are highlighted in yellow. B) Electrostatic surface representation of PSHCP shown in two orientations. The electrostatic surface was generated using APBS (42), with blue and red representing positively charged and negatively charged surfaces, respectively.

**Figure 4.**
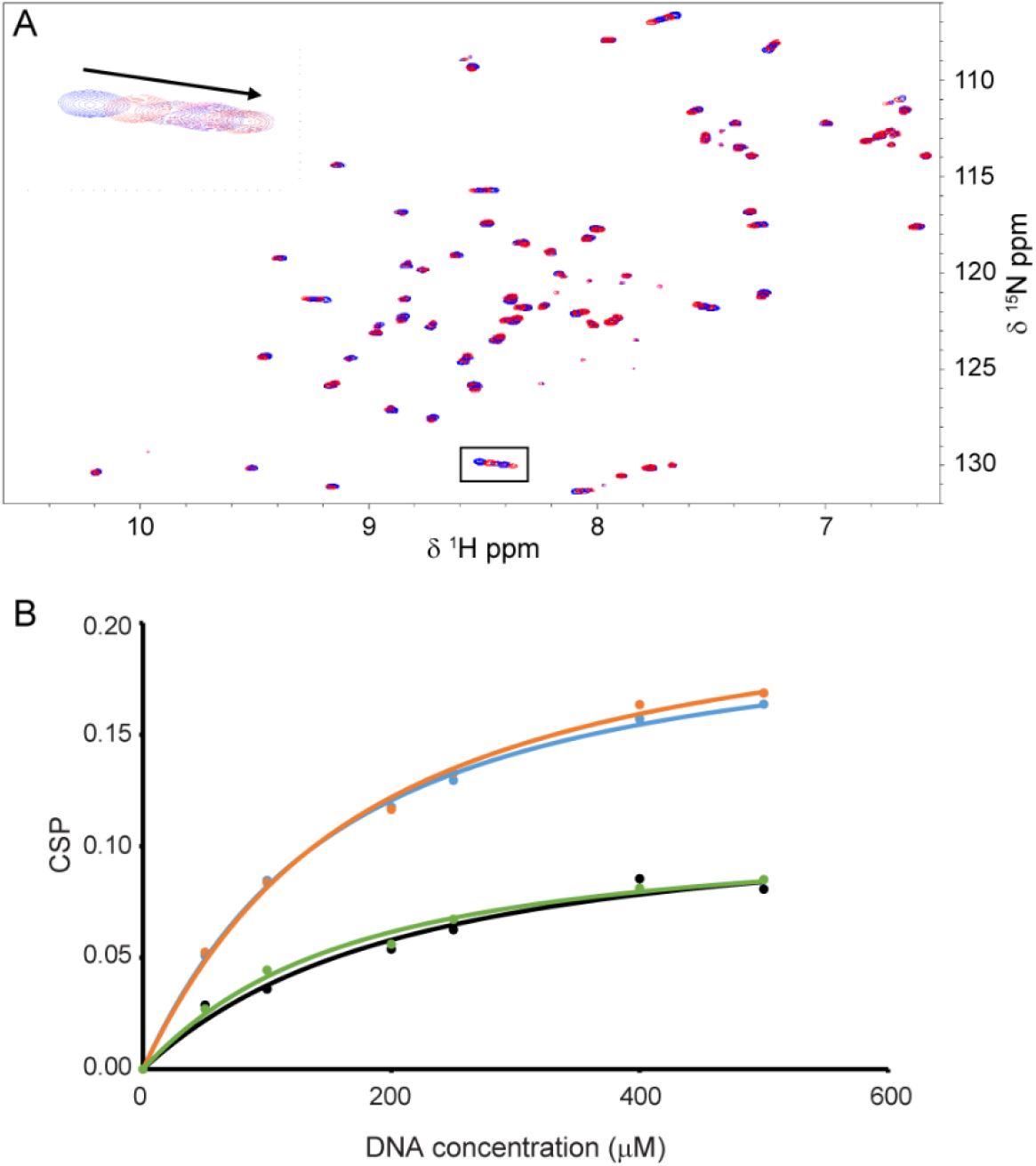
PSHCP weakly binds to dsGCDNA. A) ^1^H-^15^N HSQC spectra of 50 μM PSHCP_1-56_ in the presence of increasing concentrations of dsGCDNA. The ^1^H-^15^N HSQC spectrum of PSHCP in the absence of dsGCDNA is shown in blue. The inset shows the chemical shift perturbation for E15 with the arrow pointing in the direction of the shift with increasing DNA concentration. B) CSP data for E15 (cyan), G20 (orange), V22 (black) and T46 (green) as a function of DNA concentration. All data points are shown as circles. The fits to the data are shown as lines of the same color as the datapoints.

To determine the dissociation constant (K_d_) for PSHCP_1-56_-DNA binding, we monitored chemical shift perturbations (CSPs) at the four residues which undergo the largest CSPs in PSHCP (E15, G20, V22, T46) over a concentration range of 50 μM to 500 μM dsGCDNA (Fig. 4B and Table 2). The calculated K_d_ values ranged from 121.3 to 173.3 μM with an average K_d_ of 140.6 μM. This data demonstrates that PSHCP has a weak affinity for double-stranded DNA, which is similar to the previously measured affinities of Tudor domains for DNA. In comparison, the double interdigitated Tudor domain of the histone binding protein RBBP1 has a K_d_ of 84 μM for the same DNA sequence (23).

**Table 2.** DNA Binding Data

| Residue | Calculated $K_d$ ($\mu\text{M}$) | $R^2$ |
| --- | --- | --- |
| E15 | 121.3 | 0.99837 |
| G20 | 138.4 | 0.99680 |
| V22 | 173.3 | 0.97596 |
| T46 | 129.2 | 0.99165 |

### PSHCP binds tRNA with a low micromolar affinity

While most Tudor domains that have been studied are involved in the regulation of DNA, some Tudor domains are located in RNA-binding proteins and are involved in the regulation of RNA metabolism (21,25,26). Specifically, the knotted Tudor domain from *Saccharomyces cerevisiae* Esa1 has been shown to bind oligo-RNAs (27), and more recently the Tudor domain of *Escherichia coli* ProQ was shown to interact with the small non-coding RNA SraB (28). All of this supports a broader role for Tudor domains in the regulation of nucleic acids, including through direct binding of RNA.

Intriguingly, the PSHCP gene in completely sequenced genomes of marine *Synechococcus* and *Prochlorococcus marinus* spp. is flanked on the 5’ end by a gene encoding a tryptophan tRNA and on the 3’ end by genes encoding an aspartic acid tRNA and the glutamyl-tRNA synthetase (Fig. S3 and refs. (9,10)). Additionally, in *P. marinus* str. MED4 and MIT9313 and in *Synechococcus* sp. WH8102, the PSHCP gene is sometimes co-transcribed with the Trp-tRNA gene (10). Since genes in bacteria are often clustered and co-transcribed based on their joint functionality in a process or a pathway, we hypothesized that PSHCP may interact with tRNA. To test this, we titrated a mixture of tRNA isolated from *E. coli* into ^15^N labeled PSHCP_1-56_ and monitored CSPs in a series of ^1^H-^15^N HSQC spectra (Fig. 5A). At a 1.6:1 mixture of tRNA and PSHCP_1-56_ almost every peak in the ^1^H-^15^N HSQC had broadened to the point where it was no longer visible (Fig. S4). Only five peaks did not completely broaden during the tRNA titration and these peaks corresponded to residues at the termini of PSHCP (residues 1-4 and 56). Due to the substantial broadening of the peaks in the ^1^H-^15^N HSQC we were only able to monitor CSPs for residues 2-4 which still had visible peaks after completion of the titration. Calculated K_d_ values for PSHCP_1-56_-tRNA binding ranged from 2.9 to 10.4 μM with an average K_d_ of 7.6 μM (Fig. 5B and Table 3). As such, PSHCP displays an 18.5-fold higher affinity for tRNA than dsDNA, suggesting a potential role for PSHCP in protein translation.

**Table 3.** tRNA Binding Data

| Residue | Calculated $K_d$ ($\mu\text{M}$) | $R^2$ |
| --- | --- | --- |
| E2 | 10.4 | 0.99510 |
| L3 | 2.9 | 0.97705 |
| D4 | 9.4 | 0.98226 |

**Figure 5.**
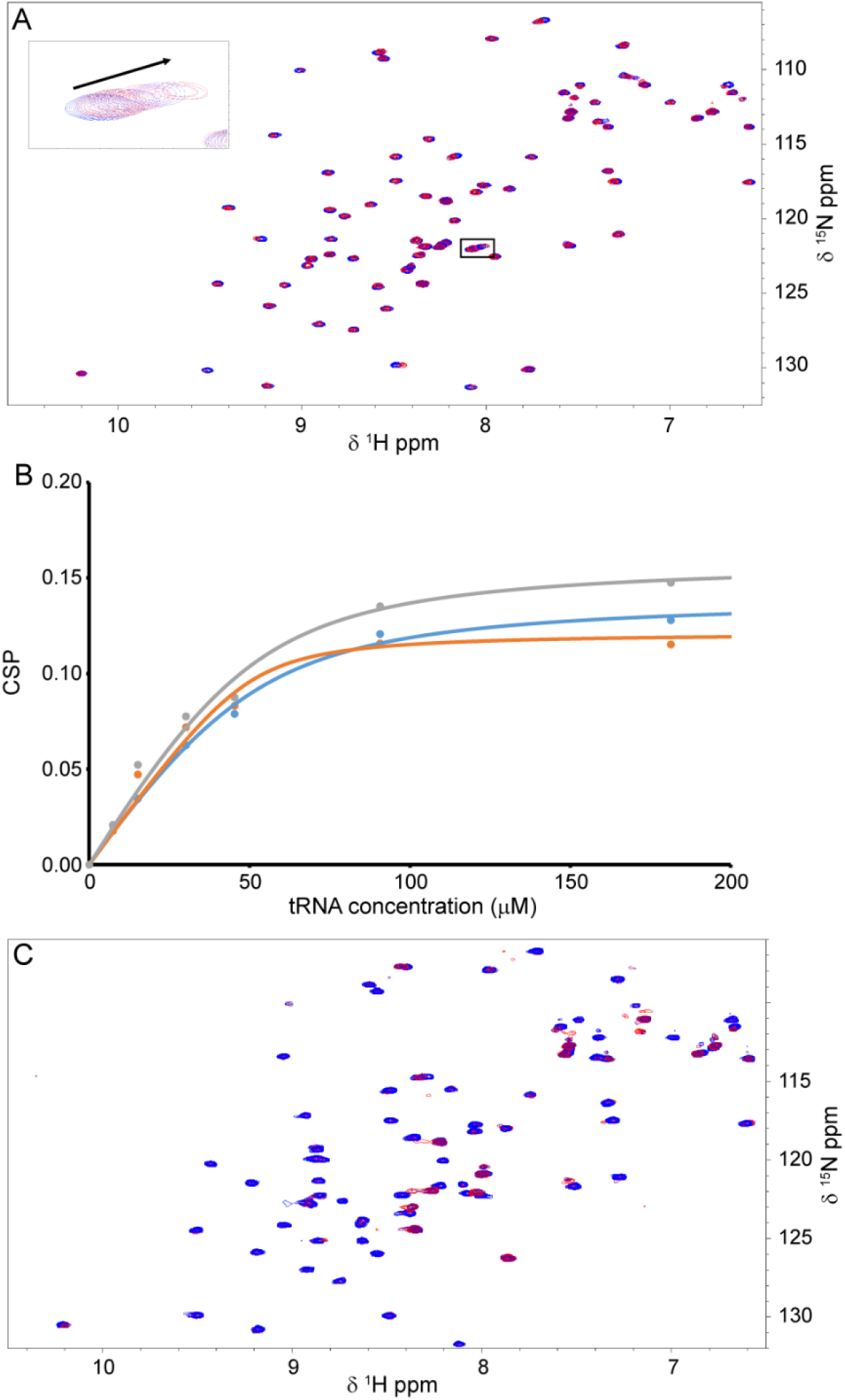
PSHCP binding to *E. coli* tRNA. A) ^1^H-^15^N HSQC spectra of 50 μM PSHCP_1-56_ with increasing concentrations of *E. coli* tRNA. The inset shows the chemical shift perturbation for L3 with the arrow showing the direction of the shift with increasing tRNA concentration. B) CSP data for E2 (cyan), L3 (orange), and D4 (grey) as a function of tRNA concentration. C) ^1^H-^15^N HSQC spectra of 50 μM PSHCP_1-62_ with increasing concentrations of *E. coli* tRNA.

To determine if the C-terminal tail of PSHCP (residues 57-62) plays a role in tRNA binding we repeated the HSQC titration experiment with ^15^N labelled PSHCP_1-62_. The addition of tRNA also led to significant line broadening of the PSHCP_1-62_ ^1^H-^15^N HSQC spectra (Fig. 5C). While peak broadening was similar between PSHCP_1-56_ and PSHCP_1-62_, clear peak shifts at the N-terminus were no longer observed. As such, we were unable to determine a K_d_ from these spectra.

### Multiple PSHCP molecules bind to a single tRNA

To determine a K_d_ for both PSHCP_1-56_ and PSHCP_1-62_ we utilized isothermal titration calorimetry (ITC) (Fig. 6). K_d_ values for PSHCP_1-56_ and PSHCP_1-62_ binding to tRNA were 1.32 μM and 1.39 μM, respectively, suggesting that residues 57-62 do not play an important role in tRNA binding (Table 4). In addition, the stoichiometry (N) measured for these binding experiments were 0.22 and 0.17 for 1-56 and 1-62, respectively. This suggests that multiple PSHCP molecules are able to bind to a single tRNA. An N around 0.2 suggests that five PSHCP molecules may bind to a single tRNA molecule. This would result in a complex of approximately 61 kDa for PSHCP_1-62_ binding to a single tRNA. This is consistent with the line broadening that was observed in the ^1^H-^15^N HSQC spectra during the titration with tRNA as line broadening can occur as a result of increasing molecular weight.

**Table 4.** ITC Data

| Sample | K<sub>d</sub> (μM) | N | ΔH (x 10<sup>4</sup> cal/mol) | ΔS (cal/mol/deg) |
| --- | --- | --- | --- | --- |
| **50 mM NaCl** |  |  |  |  |
| PSHCP<sub>1-56</sub> + tRNA | 1.32 ± 0.012 | 0.219 ± 0.142 | -5.11 ± 1.13 | -145 ± 38.2 |
| PSHCP<sub>1-62</sub> + tRNA | 1.39 ± 1.08 | 0.172 ± 0.020 | -3.60 ± 2.21 | -94 ± 72.9 |
| **25 mM NaCl 75 mM KCl** |  |  |  |  |
| PSHCP<sub>1-56</sub> + tRNA | 3.93 ± 1.21 | 0.212 ± 0.077 | -2.61 ± 0.699 | -62.7 ± 24.5 |
| PSHCP<sub>1-62</sub> + tRNA | 3.80 ± 0.673 | 0.164 ± 0.023 | -3.64 ± 0.544 | -97.8 ± 18.7 |
| PSHCP<sub>1-62</sub> + tRNA<br>Competition | 3.20 ± 0.072 | 0.162 ± 0.050 | -2.87 ± 0.305 | -71.25 ± 10.2 |
| PSHCP<sub>1-62</sub> + ssRNA | 6.74 ± 0.289 | 0.634 ± 0.182 | -2.01 ± 0.322 | -43.8 ± 10.7 |
| PSHCP<sub>1-62</sub> + dsRNA | 4.21 ± 0.656 | 0.331 ± 0.141 | -5.32 ± 3.45 | -154 ± 116 |
| PSHCP<sub>1-62</sub> + hpRNA | 5.93 ± 0.397 | 0.192 ± 0.046 | -4.74 ± 0.479 | -135 ± 16.3 |

**Figure 6.**
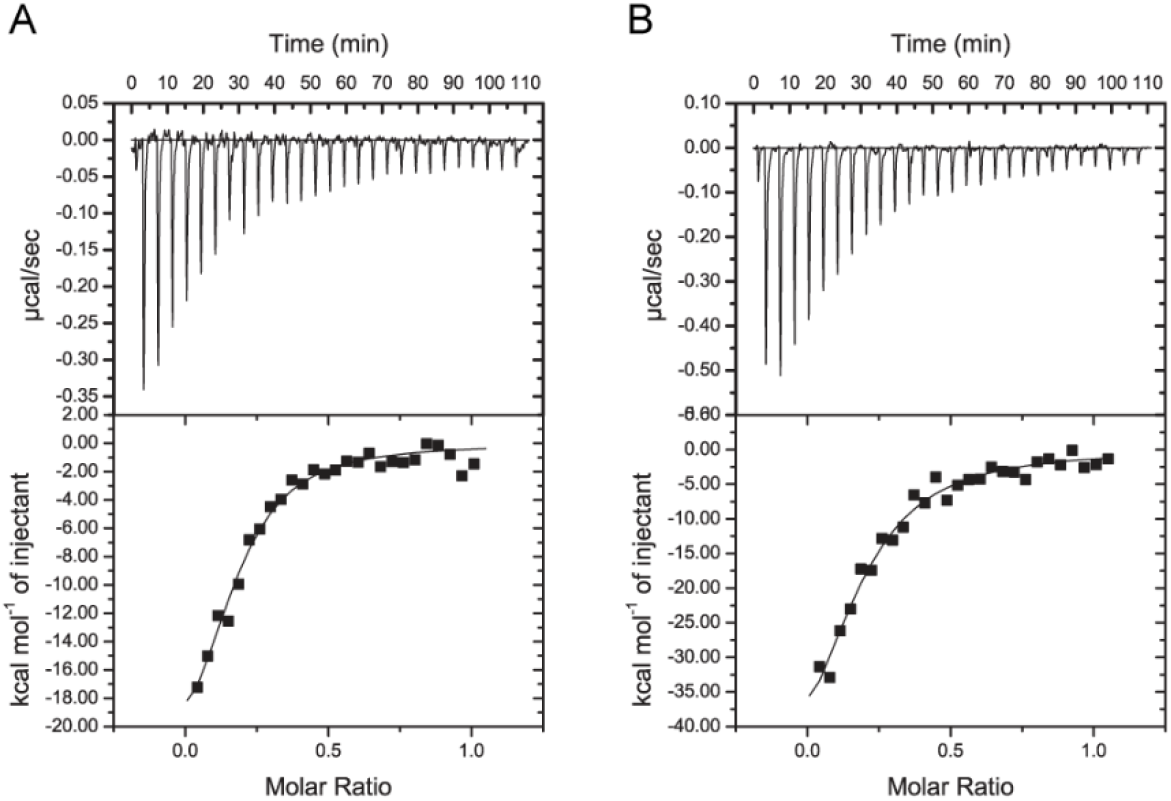
Residues 57-62 do not alter tRNA binding. A) ITC thermogram (top) and binding isotherm (bottom) for tRNA titrated into PSHCP_1-56_ in buffer containing 50 mM NaCl. B) ITC thermogram (top) and binding isotherm (bottom) for tRNA titrated into PSHCP_1-62_ in buffer containing 50 mM NaCl.

To determine if the line broadening that we observed in the ^1^H-^15^N HSQC spectra was due to the increased molecular weight from the formation of the PSHCP-tRNA complex we purified ^2^H ^15^N PSHCP_1-62_. We titrated tRNA into the ^2^H ^15^N PSHCP_1-62_ and recorded a series of ^1^H-^15^N TROSY spectra. If the line broadening in Figure 5 was due to the increased molecular weight from the formation of the PSHCP-tRNA complex, then the ^1^H-^15^N TROSY spectra should have less line broadening and allow for visualization of the peaks upon complex formation. Upon titrating tRNA into _2_H _15_N PSHCP_1-62_ we observed less line broadening in the ^1^H-^15^N TROSY spectra and clear peak shifts are now visible (Fig. 7A). This result is therefore in good agreement with a model where multiple PSHCP molecules bind to a single tRNA.

**Figure 7.**
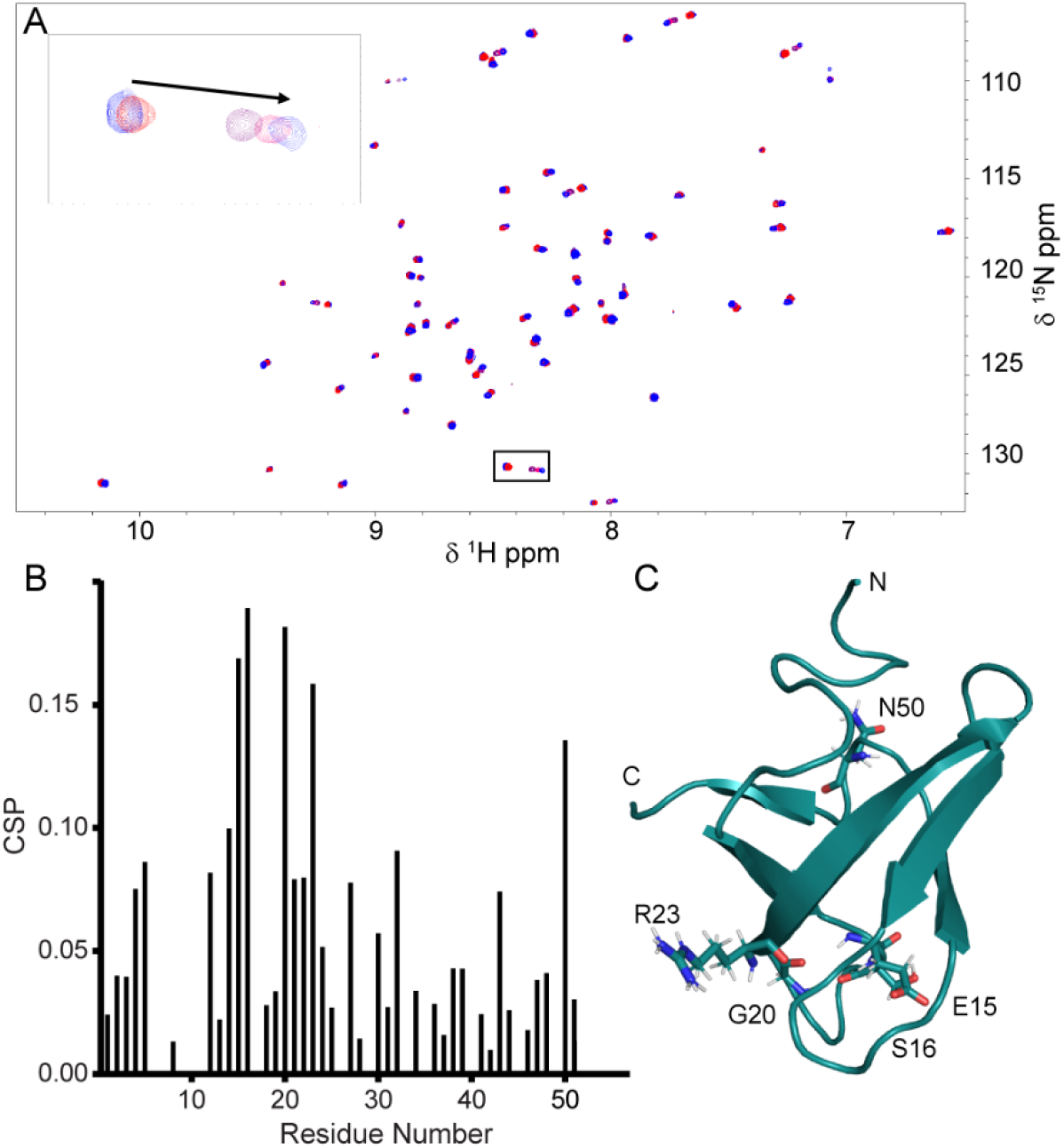
Mapping the tRNA binding interface on PSHCP. A) ^1^H-^15^N TROSY spectra of 30 μM ^2^H ^15^N PSHCP_1-62_ with increasing concentrations (0.5 to 100 μM) of *E. coli* tRNA. The inset shows the chemical shift perturbation for E15 with the arrow pointing in the direction of the shift with increasing tRNA concentration. B) CSP data for PSHCP_1-62_ in the presence of 100 μM tRNA. Peak assignments that could not be transferred from the PSHCP_1-56_ assignment were given no CSP value. C) Cartoon representation of PSHCP showing the five residues with the highest CSPs as stick representations. Each of the five amino acids is labeled along with the N and C termini.

### Residues involved in tRNA binding

To determine which regions of PSHCP bind to tRNA, CSPs were monitored in the ^1^H-^15^N TROSY spectra for PSHCP_1-62_ in the presence of 100 μM tRNA (Fig. 7B). The five largest CSPs observed were for residues E15, S16, G20, R23 and N50 (Fig. 7C). Residues E15, S16 and G20, all of which sit in the loop between β-strands 1 and 2, also shifted upon DNA binding suggesting that PSHCP utilizes similar interaction surfaces for both DNA and tRNA binding.

Since our NMR experiments were performed at 50 mM NaCl we wanted to determine if PSHCP binds tRNA at higher salt concentrations which may more closely mimic physiological salt concentrations. Intracellular NaCl concentrations have not been measured for marine *Prochlorococcus* and *Synechococcus* which live in environments with high NaCl concentrations (450-700 mM NaCl). However, measurements on a variety of other cyanobacteria species revealed that these species have intracellular Na^+^ concentrations ranging from 10-80 mM when extracellular Na^+^ concentrations are 300-500 mM (29). In addition, higher K^+^ concentrations or other compatible solutes like sucrose are frequently utilized to prevent hypertonic stress (30). As such, we monitored tRNA binding to PSHCP in a buffer containing a mixture of 25 mM NaCl and 75 mM KCl using ITC (Fig. 8A and 8B). The K_d_ of PSHCP for tRNA was slightly reduced in this buffer to 3.93 μM and 3.80 μM for PSHCP_1-56_ and PSHCP_1-62_, respectively (Table 4).

**Figure 8.**
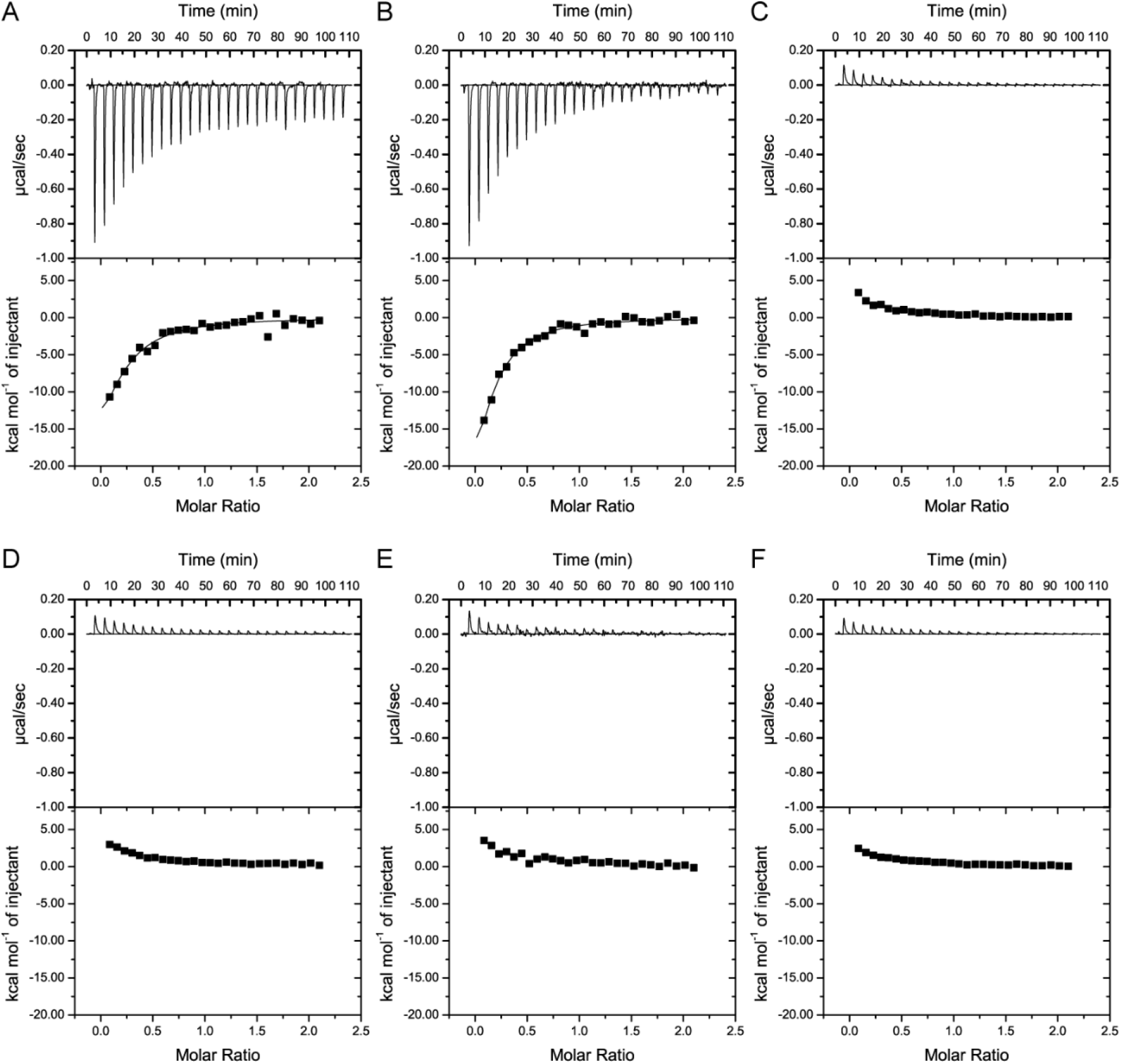
Binding site mutations on PSHCP. ITC was performed titrating tRNA into different PSHCP samples in buffer containing 25 mM NaCl and 75 mM KCl. ITC thermogram (top) and binding isotherm (bottom) for A) tRNA titrated into PSHCP_1-56_, B) tRNA titrated into PSHCP_1-62_, C) tRNA titrated into PSHCP_1-62_ E15R23AA, D) tRNA titrated into PSHCP_1-62_ E15K30AA, E) tRNA titrated into PSHCP_1-62_ S16K30AA, F) tRNA titrated into buffer alone.

PSHCP contains two positively charged patches on its surface (Fig. 3). Our CSP data suggest that the surface surrounding R23, including β-strand 2 and the loop in between β-strand 1 and 2, is the main interaction site for tRNA. To further test this we chose to mutate E15, S16 and R23 to alanine as these residues had the largest CSPs during the tRNA titration (Fig. 7). We also chose to mutate K30 to alanine as this residue is located in the other positively charged patch located on the opposite side of PSHCP from R23. We generated three double mutants in PSHCP_1-62_, E15R23AA, E15K30AA and S16K30AA and performed ITC experiments in buffer containing 25 mM NaCl and 75 mM KCl (Fig. 8 C-E). The exothermic peaks that were previously observed for PSHCP binding to tRNA were not visible in the titrations with any of the mutant proteins suggesting that none of these proteins bind tRNA. Small endothermic peaks are visible, but these peaks are also observed when tRNA is titrated into buffer alone demonstrating that these peaks are the result of tRNA dilution and not a binding interaction (Fig. 8F). Taken together, our CSP and mutagenesis data show that E15, S16 and R23 are all important for tRNA binding.

### PSHCP has a broad specificity for RNA

To determine if PSHCP binding is specific for tRNA, we initially performed competition binding experiments using ITC. In these experiments, PSHCP_1-62_ was incubated with an equimolar concentration of dsGCDNA in buffer containing 25 mM NaCl and 75 mM KCl for 30 minutes. ITC was then performed with tRNA being titrated into the mixture of PSHCP_1-62_ and dsGCDNA. In these experiments a K_d_ of 3.20 μM and an N of 0.162 was observed which is consistent with the interaction of PSHCP_1-62_ for tRNA (Fig. 9A). This demonstrates that dsGCDNA does not compete with tRNA for PSHCP binding.

**Figure 9.**
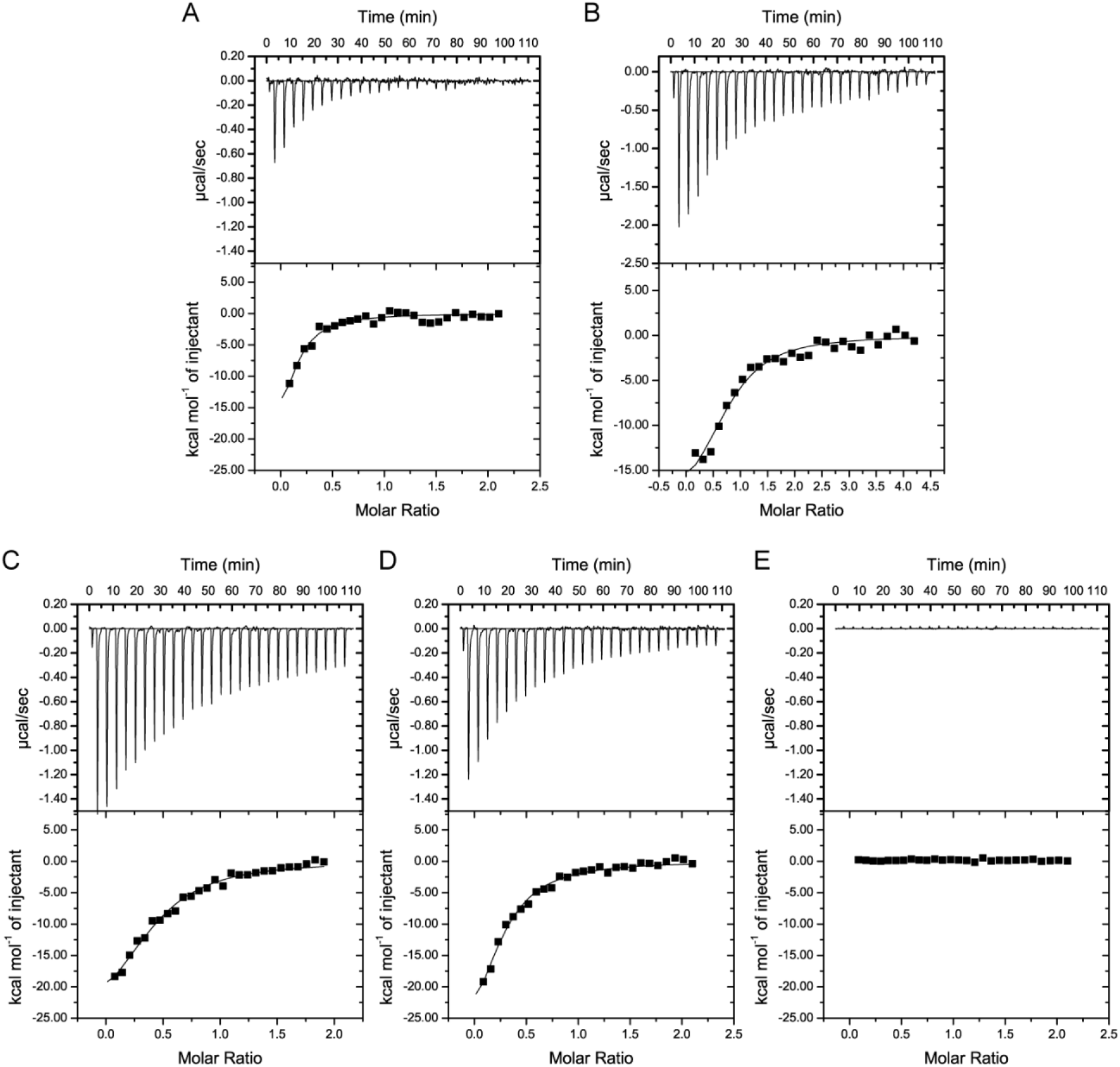
RNA binding preferences of PSHCP. ITC experiments with different RNAs being titrated into PSHCP_1-62_ in buffer containing 25 mM NaCl and 75 mM KCl. ITC thermogram (top) and binding isotherm (bottom) for A) tRNA titrated into a 1:1 mixture of PSHCP_1-62_ and dsGCDNA, B) ssRNA titrated into PSHCP_1-62_, C) dsRNA titrated into different PSHCP_1-62_, D) hpRNA titrated into PSHCP_1-62_, E) a mixture of ATP, GTP, UTP and CTP titrated into PSHCP_1-62_.

To determine which structural features of the tRNA are recognized by PSHCP we generated three different RNA fragments. These fragments were single-stranded RNA (ssRNA), double-stranded RNA (dsRNA) and an RNA hairpin (hpRNA). These sequences were derived from different regions of the aspartic acid tRNA sequence which is adjacent to the PSHCP gene. PSHCP was able to bind to all three of these RNAs with K_d_ values ranging from 4.21 to 6.74 μM suggesting that PSHCP has a broad specificity for RNA (Fig. 9B-D and Table 4). In addition, the N for dsRNA and hpRNA was similar to the N for tRNA, suggesting that multiple PSHCP molecules can bind to a single dsRNA or hpRNA. In contrast, the N for PSHCP_1-62_ binding to ssRNA was 0.634 suggesting that less PSHCP molecules bind to a single ssRNA. Since PSHCP binds to dsRNA, hpRNA and ssRNA with similar affinities this suggests that PSHCP is likely recognizing the phosphodiester backbone of RNA and not any specific sequence or structure. To further test this, we performed ITC with a mixture of ATP, GTP, UTP and CTP (Fig. 9E). We found that PSHCP was unable to bind to these individual nucleotides. Taken together, our data demonstrate that PSHCP has a broad specificity for RNA.

## DISCUSSION

We used NMR spectroscopy to determine the structure of the highly conserved 62 amino acid cyanobacterial protein PSHCP. PSHCP residues 4-56 contain a Tudor domain, with five β-strands arranged in a barrel-like architecture. The remainder of the protein is disordered. Two positively charged patches on the surface of PSHCP hinted that it may bind directly to nucleic acids. We found that PSHCP binds to a variety of RNAs with low micromolar affinities. This suggests that PSHCP may likely be involved in RNA regulation. We also found that residues 57-62 did not enhance the binding affinity of PSHCP for tRNA and it remains to be seen what role this disordered tail may have in PSHCP function.

The majority of the Tudor domains that have been studied are eukaryotic Tudor domains that regulate histone function by interacting with methylated lysine residues (20). However, there are several examples of Tudor domains that bind directly to nucleic acids instead of methylated proteins. These nucleic acid binding Tudor domains are broadly distributed from bacteria to eukaryotes, suggesting that the ancestral function of Tudor domains may be nucleic acid binding rather than protein binding. Structures of such nucleic acid binding Tudor domains have been determined for the DNA damage response protein 53BP1 from *Homo sapiens* (31), the histone binding protein RBBP1 from *H. sapiens* (23), the methyltransferase Esa1 from *S. cerevisiae* (27), and the RNA binding protein ProQ from *E. coli* (28). The Tudor domain of PSHCP shares structural similarity with the core region of these four nucleic acid binding Tudor domains (Fig. S5), but there is a surprising divergence in the mechanism of nucleic acid binding between these different Tudor domains, as detailed below.

Both 53BP1 and RBBP1 contain two adjacent Tudor domains (23,31). 53BP1 contains two sequential Tudor domains that are separated in primary sequence and do not share any secondary structure (Fig. S5A). In contrast, RBBP1 forms an interdigitated Tudor domain where some of the secondary structure of the two Tudor domains alternate in the primary sequence (Fig. S5B). In addition, the two Tudor domains of RBBP1 share 2 β-strands generating a larger rigid scaffold for DNA binding. DNA binding to either 53BP1 or RBBP1 relies on residues in both Tudor domains (31). RBBP1 contains a positively charged pocket in between the two Tudor domains that allows for DNA binding (23). This is a distinct mode of nucleic acid binding formed by the overall interdigitated structure of the two Tudor domains. PSHCP aligns well to the individual Tudor domains from these structures but does not contain an extended surface like that of 53BP1 or RBBP1 for nucleic acid binding.

Esa1 contains a knotted Tudor domain due to the presence of two additional β-strands (β0 and β6) which lie just before and just after the classical Tudor domain (Fig. S5C) (27). PSHCP aligns well to the short version of the Esa1 Tudor domain, which lacks these additional β-strands. The short version of Esa1 is incapable of binding to RNA while the knotted Tudor domains binds to poly(U) RNA with a 21.6 μM K_d_ in buffer containing 10 mM NaCl (27). This binding interaction has the highest affinity of any of the interactions tested. It is surprising then that PSHCP is able to bind RNA with a low micromolar affinity even though it is most similar in structure to the short version of Esa1 which is incapable of binding RNA.

The *E. coli* ProQ protein contains two domains, an N-terminal FinO like domain and a C-terminal Tudor domain (28). Both of the domains interact with small RNAs to facilitate binding with low nanomolar affinities (32). The primary RNA interaction region on ProQ appears to be through the FinO domain (28). However, two regions on the ProQ Tudor domain were also shown to interact with RNA. One of these regions spans β-strands 2 and 3. This region has some overlap with the primary RNA binding surface on PSHCP which also includes β-strand 2. While these Tudor domains have some common RNA binding surfaces, the location of the positively charged amino acids in these regions are in different β-strands. In PSHCP, R23 is located on β-strand 2 while in ProQ R214 is located on β-strand 3. The binding affinity of the ProQ Tudor domain for RNA has not been experimentally determined and therefore it is unclear if the ProQ Tudor domain has a similar affinity for RNA as PSHCP.

All of the discussed ‘nucleic acid binding’ Tudor domains exist as part of multidomain proteins that range in size from 232 to 1257 amino acids. The interaction of these proteins with nucleic acids often requires more than one domain for efficient binding. Since PSHCP is only 62 amino acids, it is rather remarkable that it has such a strong binding affinity for a variety of RNA molecules in such a small number of amino acids. It will be interesting to determine exactly what role the PSHCP-RNA interaction plays in a subgroup of *Synechococcales* and why it contains such a broad specificity for RNA. Does this broad specificity allow PSHCP to bind to a large number of different RNA targets in cells or is there a specific target that has not been identified? In pull-down assays, PSHCP was observed to be associated with the ribosomal protein L2 (10). Taken together with its RNA binding ability reported here, it is tempting to speculate that PSHCP is part of the ribosome complex or perhaps is involved in protein translation. Such functions could explain the extraordinary amino acid conservation of the protein, as ribosomal and translational machinery proteins are among the most conserved. Further studies are needed to evaluate these hypotheses.

## EXPERIMENTAL PROCEDURES

### Protein Expression and Purification

PSHCP_1-56_ and PSHCP_1-62_ were cloned into the ligation independent cloning vector (1B) which was a gift from Scott Gradia (Addgene, 29653). Plasmids containing PSHCP were transformed into BL21 (DE3) star cells (Invitrogen, C601003). PSHCP mutants were generated using Q5 Site-Directed Mutagenesis Kit (NEB, E0554S). Cells were grown in Terrific Broth (Fisher, BP9728-2) to an optical density at 600 nm of approximately 3.0. For unlabeled protein expression was induced with 1 mM isopropylthio-β-D-galactoside (IPTG; Amresco, 97061) and cultures were grown for 18 hours at 18°C. For uniformly labelled ^15^N or ^15^N and ^13^C protein cells were pelleted (3,000 x g, 20 min) and resuspended in M9 minimal media comprising 3 g/L ^15^N ammonium chloride (Cambridge Isotope Laboratories, NLM-467-25) with 10 g/L glucose or 10 g/L ^13^C glucose (Cambridge Isotope Laboratories, CLM-1396-25). Expression was induced using 1 mM isopropylthio-β-D-galactoside (IPTG; Amresco, 97061) and cultures were grown for 18 hours at 18°C. Cells were harvested and stored at −80°C. Cell pellets were thawed and resuspended in 50 mM Tris, pH 8.0, 500 mM NaCl, 5 mM MgCl_2_, 0.1% (v:v) Triton X-100 buffer containing protease inhibitors (Roche, 11836170001). Cells were lysed using a French Press (Thermo Electron, FA-032) and lysates were cleared by centrifugation for 50 min at 40,000 xg at 4°C. Cleared lysates were incubated with TALON resin (Clontech, 635504) which was preequilibrated with 50 mM Tris, pH 8.0, 500 mM NaCl. Protein was eluted in buffer containing 200 mM imidazole. TEV protease was used to cleave off the hexahistidine tag. Cleaved protein was flowed over a desalting column to remove imidazole and a second purification using TALON resin was used to remove the TEV protease and any uncleaved protein. A final purification step was carried out using a HiLoad Superdex 75 PG column (GE Healthcare, 28-9893-33) equilibrated in 20 mM sodium phosphate pH 6.5 and 50mM NaCl or 25mM NaCl, 75mM KCl for nucleotide binding experiments or 20 mM sodium phosphate pH 6.5, 200 mM NaCl, and 0.2 mM tris(2-carboxyethyl)phosphine (Amresco, K831-26) for structural studies.

### Deuterated Protein Expression and Purification

PSHCP_1-62_ (1B) was transformed into BL21 RILP cells (Agilent, 230280). For the production of deuterated, uniformly ^15^N-labelled protein, we followed the protocol outlined in Tugarinov et.al. (33). Briefly, cells were grown in the presence of ^15^N ammonium chloride (Cambridge Isotope Laboratories, NLM-467-25), deuterated glucose (Cambridge Isotope Laboratories, DLM-2062-10), and deuterium oxide (Cambridge Isotope Laboratories, DLM-4-1000). Protein expression was induced with 100 μM IPTG and cultures were grown for 18 hours at 18°C. Protein was purified as in *Protein Expression and Purification* in 20mM sodium phosphate pH 6.5 and 50mM NaCl.

### RNA Synthesis

RNA was produced by *in vitro transcription* (IVT) using the HiScribe T7 High Yield RNA Synthesis Kit (New England BioLabs, E2040S). DNA templates for the IVT reaction were produced by annealing complementary DNA oligos (IDT) for the T7 polymerase promoter sequence (TAATACGACTCACTATAGGG) and the appropriate RNA template. Oligos were resuspended in Annealing Buffer (10mM Tris pH 8.0, 50mM NaCl) and mixed in equimolar amounts, heated at 94°C for 2 minutes, then cooled to room temperature to achieve annealing of the two oligos. IVT reactions products were purified using the Monarch RNA Cleanup Kit (New England BioLabs, T2050S). RNA products were analyzed by A260/A280 for purity using a NanoDrop instrument.

RNAs produced were ssRNA (GGGUUGAACUGGUUA), dsRNA (GGGUAACCAGUUCAA annealed to ssRNA), and hpRNA (GGGCGCCUGUCACGGCG).

### NMR Spectroscopy and Structure Determination

All NMR experiments for assignment and structure determination were performed at 298 K on a 700 MHz Bruker Avance III spectrometer. The sequence-specific backbone assignment was determined using 2D [^1^H-^15^N] HSQC, 3D HNCA, 3D HN(CO)CA, 3D HNCO, 3D HN(CA)CO, 3D CBCA(CO)NH, 3D HNCACB and 3D HBHA(CBCACO)NH experiments. Aliphatic side chain assignments were determined using 3D (H)CCH-TOCSY, 3D HC(C)H-COSY, 3D HC(C)H-TOCSY, 3D H(CCCO)NH and 3D (H)CC(CO)NH experiments. Amide side chain assignments were determined using a 3D ^15^N-resolved NOESY. ^1^H chemical shifts were externally referenced to 0 ppm methyl resonance of 2,2-dimethyl-2-silapentane-5-sulfonate (DSS), whereas ^13^C and ^15^N chemical shifts were indirectly referenced according to the IUPAC recommendations (34). All NMR spectra were processed using Topspin 3.5 (Bruker). Processed spectra were analyzed using CARA (http://cara.nmr.ch/). Backbone and side chain assignments were deposited in the BMRB under accession 30559.

For structure calculation a 2D ^1^H-^1^H NOESY, 3D ^15^N-resolved NOESY and 3D ^13^C-resolved NOESY were recorded and processed using Topspin 3.5 (Bruker). UNIO was used for iterative automated NOE peak picking and NOE assignment by ATNOS/CANDID (35,36) and structure calculation with CYANA v2.1 (37,38). The 20 lowest energy structures were used for water refinement in CNS v1.3 with the RECOORD scripts (39). The quality of the final ensemble was verified using NMR-Procheck and the RMSD of these structures was determined using MolMol (40,41). The final structure was deposited in the PDB under ID 6NNB. All structure figures were generated using the PyMOL Molecular Graphics System, Version 1.8 (Schrödinger, LLC). APBS was used to generate the electrostatic surface representation of PSHCP (42). To determine the intensity ratio of peaks from the {^1^H}-^15^N heteronuclear NOE experiment, peak intensity values were determined using CARA (http://cara.nmr.ch/). Experimental uncertainties for the {^1^H}-^15^N heteronuclear NOE experiment were determined by measuring the root-mean-square (RMS) noise of background regions in the unsaturated and saturated spectra using NMRPipe (43) as described previously (44).

### NMR Titration Experiments

dsGCDNA (5′-CCG CGC GCG CGG-3′) was synthesized by IDT. DNA was resuspended in 20mM sodium phosphate pH 6.5, 50 mM NaCl, heated at 100°C for 5 min and cooled at room temperature to allow for the DNA to anneal. tRNA from *E. coli* (Sigma R1753) was resuspended in 20 mM sodium phosphate pH 6.5, 50mM NaCl. The concentration of the *E. coli* tRNA was determined by running a range of concentrations of the tRNA on an agarose gel and comparing them to a standard concentration. ^15^N labeled PSHCP was mixed with nucleic acid samples to a final protein concentration of 50 μM. The DNA and tRNA concentrations were varied based on the overall titration. ^1^H-^15^N HSQC spectra were recorded on a Bruker Avance 600 MHz spectrometer using a 1.7 mm probe. For the TROSY titrations, ^2^H, ^15^N labeled PSHCP_1-62_ was mixed with tRNA and ^1^H-^15^N TROSY spectra were collected on a 700 MHz Bruker Avance III spectrometer. For these titrations, PSHCP_1-62_ was at a final concentration of 30 μM and tRNA was at a final concentration of 0.5, 25, 50 or 100 μM. CSP values were determined using δΔ = √((δ^1^H)_2_ + 0.14(δ^15^N)_2_). CSP data were fit using MATLAB (MathWorks) to f(x) = CSP_max_*((K_d_+x+P_tot_)-sqrt((K_d_+x+P_tot_)^2-4*(x*P_tot_)))/(2*(P_tot_)) where P_tot_ is the total concentration of protein (50 μM).

### Isothermal titration calorimetry

ITC was performed using a VP-ITC (Microcal). All proteins and tRNA were prepared in 20mM sodium phosphate pH 6.5, 50 mM NaCl or 20mM sodium phosphate pH 6.5, 25 mM NaCl, 75 mM KCl. Approximately 200 μM *E. coli* tRNA (Sigma R1753) was titrated into 20 μM PSHCP (either PSHCP_1-62_ or PSHCP_1-56_) at 24°C. tRNA concentration was determined by running a tRNA titration on a 1% agarose gel. tRNA samples were then compared to samples of known concentration. ssRNA, dsRNA, and hpRNA were produced in-house (see *RNA Synthesis)* in the same buffer. These RNAs, at 200 μM, were titrated into PSHCP_1-62_ at 20 μM except for the ssRNA which was at 400 μM. For ITC competition assays, 20μM dsGCDNA was mixed with 20μM PSHCP_1-62_ for 30 minutes. Subsequently, 200μM tRNA was titrated into the mixture of PSHCP_1-62_ and dsGCDNA.

Data were analyzed using Origin (Microcal) to produce both ITC thermograms and binding isotherms.

### Survey of PSHCP Taxonomic Distribution and Amino Acid Conservation

The amino acid sequence of the PSHCP gene from *Prochlorococcus marinus* str. MIT 9312 (RefSeq ID WP_002807701.1) was used as a query in a BLASTP search of the *nr* database via NCBI web site (accessed on September 12, 2018; BLAST v. 2.8.0+ (45); E-value cutoff 10^−4^). Taxonomic distribution was examined via Taxonomy Report provided with the BLAST search results. Matches with sequences not identical to the query were retrieved and aligned in ClustalX v. 2.1 (46). Additionally, the same query sequence was used in a BLASTP search of the 703 genomes of *Prochlorococcus* and marine *Synechococcus* available via IMG/ProPortal (47). This database includes both isolates and single cell genomes, the latter ones with an average completion of 60% (47). The BLASTP 2.6.0+ search was carried out via IMG/MER web site at https://img.jgi.doe.gov/ on September 11, 2018, with E-value cutoff of 10^−5^. The obtained 463 PSHCP homologs were retrieved and aligned in ClustalX v. 2.1 (46). Two poor quality sequences (defined as those containing at least one X) and one partial sequence at the beginning of a short contig in an incomplete genome were removed from the alignments. Both alignments are available as Supplemental Datasets 1 and 2.

## Acknowledgments

This work was supported by Dartmouth start-up funds and the National Institutes of Health awards GM113132 and GM128663 to M.J.R, and by the Simons Foundation Investigator in Mathematical Modeling of Living Systems award 327936 to O.Z. We would like to thank Amanda Cockshutt and Kaitlyn Connelly for helpful discussions.

## Author Contributions

K.M.B. and R.D. purified protein, analyzed NMR data, and collected ITC data. K.M.B. synthesized RNA. M.P. collected all of the NMR spectra. O.Z. carried out the survey of PSHCP taxonomic distribution. O.Z. and M.J.R. conceptualized the project. K.M.B., M.P., O.Z. and M.J.R. wrote and edited the manuscript.

