## Supplemental Material for "The structure of a highly conserved picocyanobacterial protein reveals a Tudor domain with an RNA binding function"

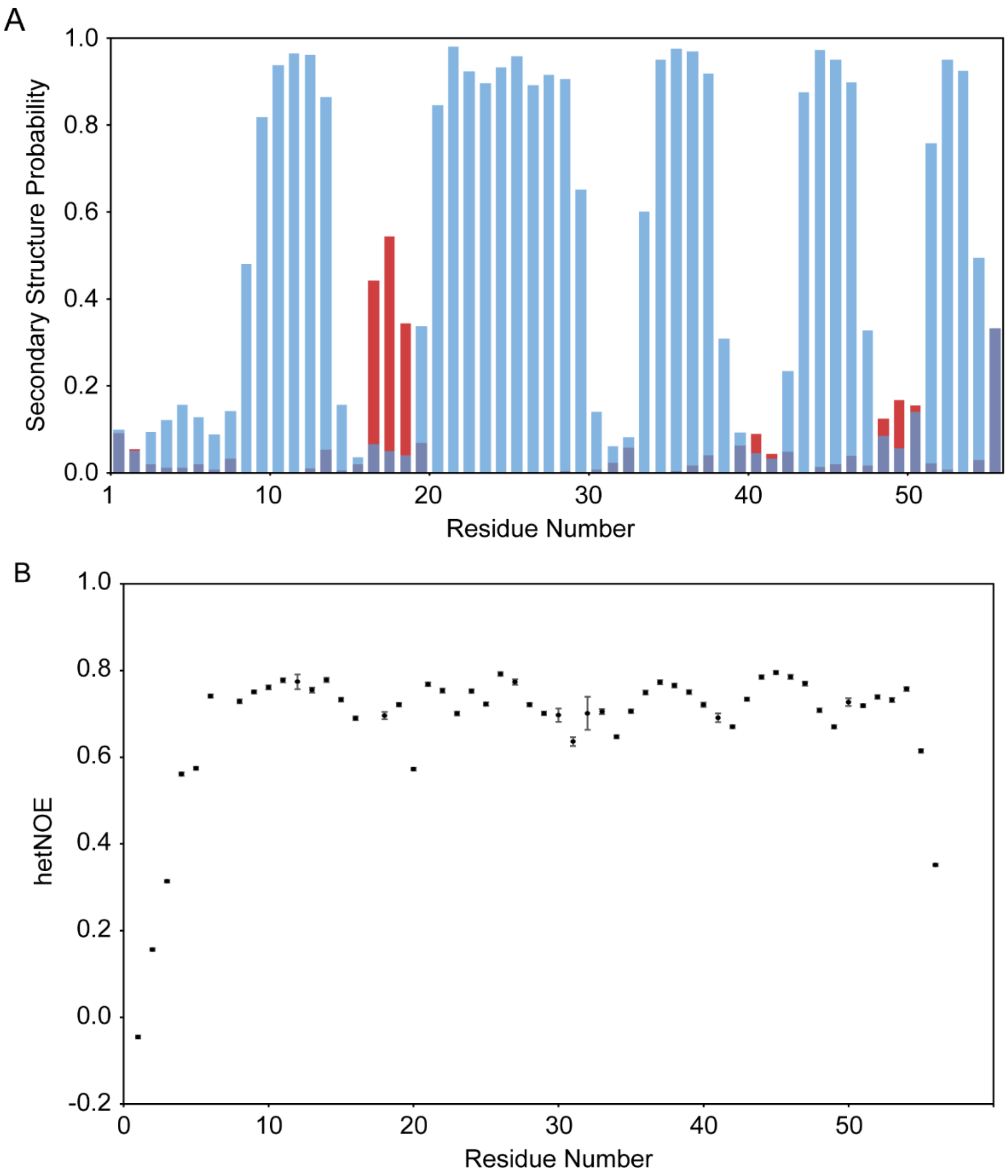


**Figure S1. Secondary structure and dynamics in PSHCP$_{1\text{-}56}$.** A) Secondary structure probabilities were calculated using the TALOS-N webserver (17). Blue bars: probability for β-strands; red bars: probability for α-helices. B) Heteronuclear $^{1}$H-$^{15}$N NOE data (hetNOE), collected for PSHCP$_{1\text{-}56}$, is plotted as the ratio of saturated/unsaturated peak intensity for each protein reside. Error bars represent the experimental uncertainties.

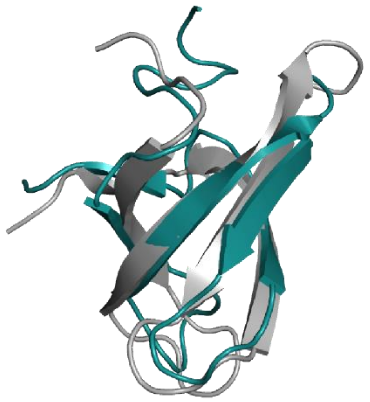

**Figure S2. Structural comparison between PSHCP and the Tudor domain of SMN.** PSHCP and the Tudor domain of SMN (PDB ID: 1G5V) are both shown as cartoon representations with PSHCP colored in cyan and SMN in grey.

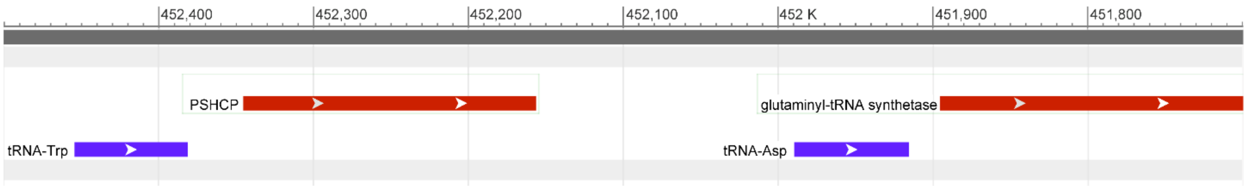


**Figure S3. Gene neighborhood of *PSHCP* in the *Prochlorococcus marinus* CCMP1375 genome.** An 800 base pair region of the *Prochlorococcus marinus* CCMP1375 genome is shown. Open reading frames coding for proteins and RNA are colored in red and purple, respectively. This figure was generated using the NCBI Graphical Sequence Viewer.

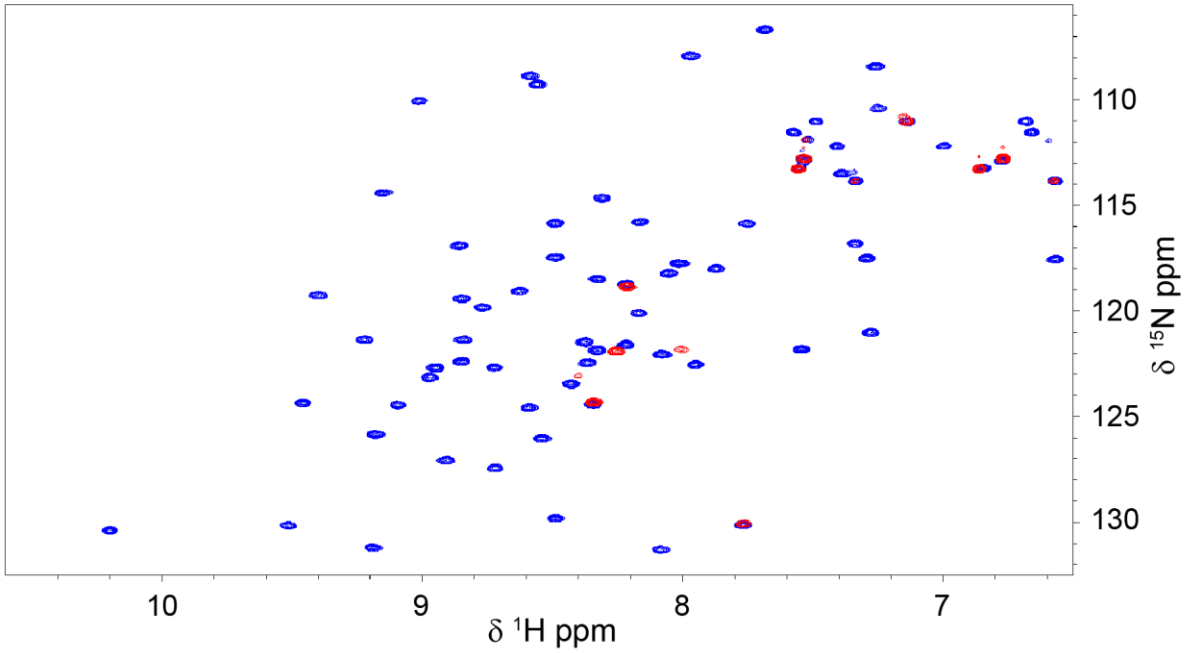


**Figure S4. Line broadening from the addition of tRNA to PSHCP.** $^{1}$H-$^{15}$N HSQC spectra of 50 µM PSHCP$_{1\text{-}56}$ in the absence of *E. coli* tRNA (blue) and in the presence of 90.7 µM *E. coli* tRNA (red).

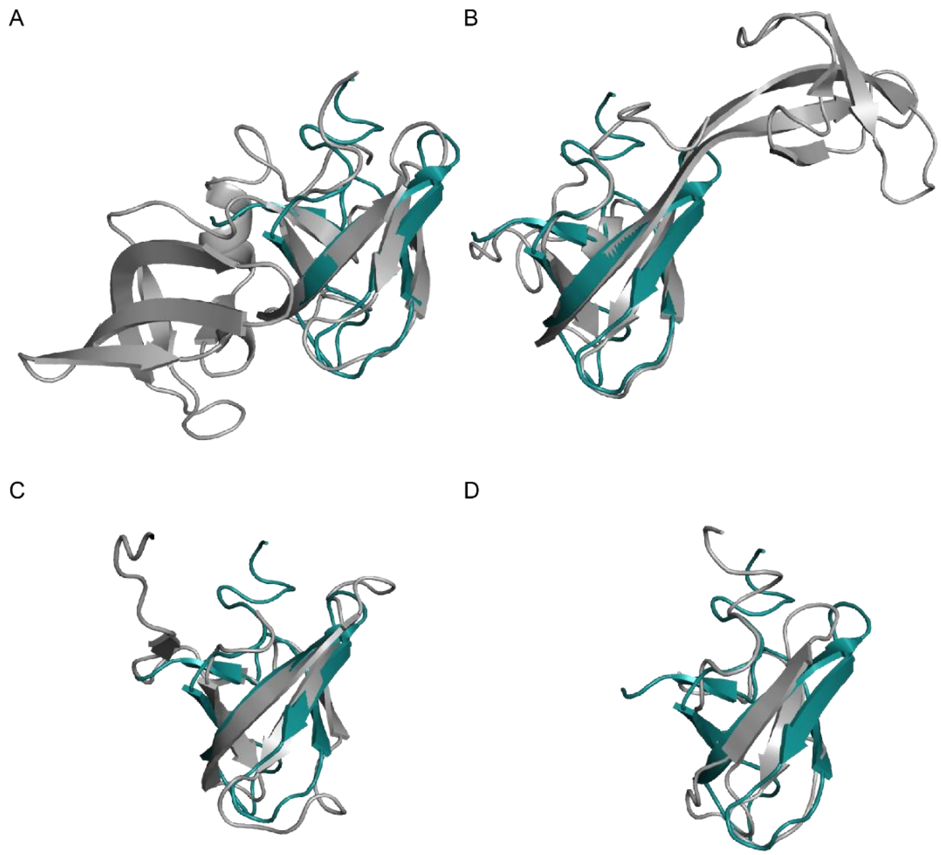


**Figure S5. Structural comparison between PSHCP and other nucleic acid binding Tudor domains.** PSHCP is shown as a cartoon representation in cyan in all panels. A) Overlay of PSHCP with the tandem Tudor domains from 53BP1 (PDB ID: 1SSF) shown in grey. B) Overlay of PSHCP with the interdigitated Tudor domain from RBBP1 (PDB ID: 2MAM) shown in grey. C) Overlay of PSHCP with the knotted Tudor domain from Esa1 (PDB ID: 2RO0) shown in grey. D) Overlay of PSHCP with the Tudor domain from ProQ (PDB ID: 1SSF) shown in grey.

**Supplementary Datasets**

**Supplementary Dataset 1.** The alignment of PSHCP homologs retrieved from the IMG/ProPortal database. The sequence IDs are IMG Gene IDs. The alignment is in FASTA format. [Filename is 'ProPortal_PSHCP_homologs_edited.fasta']

**Supplementary Dataset 2.** The alignment of PSHCP homologs retrieved from the NCBI's *nr* database. The sequence IDs The sequences were renamed according to their NCBI taxonomic classification. When strain name is not given, the record represents a MULTISPECIES record. The alignment is in FASTA format. [Filename is 'nr_non_identical_PSHCP_homologs_edited.fasta']
